# Oxytocin facilitates self-serving rather than altruistic tendencies in competitive social interactions via orbitofrontal cortex

**DOI:** 10.1101/501171

**Authors:** Xiaolei Xu, Congcong Liu, Xinqi Zhou, Yuanshu Chen, Zhao Gao, Feng Zhou, Juan Kou, Benjamin Becker, Keith M Kendrick

## Abstract

While the neuropeptide oxytocin can facilitate empathy and altruistic behavior it may also promote self-serving tendencies in some contexts, and it remains unclear if it would increase altruistic or self-interest behaviors when they compete within a social situation. The current between-subject, double-blind, placebo-controlled fMRI study investigated the effect of intranasal oxytocin on empathy for social exclusion using a modified online ball-tossing game which incorporated monetary rewards and both the potential to display altruistic and self-interest behaviors. Results showed that when subjects in both oxytocin and placebo groups were observing a player being excluded (victim) by other players in the game there was activation in the mentalizing network. When subjects then played both with the victim and the players who had excluded them they threw more balls to the victim player, indicative of an altruistic response. However, subjects in the oxytocin group threw more balls to the excluder players indicative of greater self-interest, since the latter would be perceived as more likely to reciprocate to maximize financial gain. This behavioral effect of oxytocin was associated with greater medial orbitofrontal cortex activation when playing with the excluders and negatively correlated with trait-altruism scores. Overall, our findings suggest that in the context of competing motivations for exhibiting altruistic or self-interest behavior oxytocin enhanced self-interest and this was associated with greater activation in frontal reward areas.

## Introduction

Empathy represents a core social function that allows individuals to recognize and understand the emotional states of others and respond to them accordingly (Eisenberg, & Eggum, 2009). Empathy has two main components, cognitive empathy which includes cognitive processes of perspective-taking allowing us to infer the mental states of others and emotional empathy reflecting a direct affective reaction involving understanding, sharing and responding appropriately to their feelings (Bernhardt, & Singer, 2012; Shamay-Tsoory, 2011; Shamay-Tsoory et al., 2009). Emotional empathy can further be sub-divided in direct and indirect components which have also been described as emotional contagion and a more general form of emotional arousal (Decety, 2011; Hurlemann et al., 2010). Although the cognitive and emotional components of empathy are partly dissociable (Shamay-Tsoory et al., 2009), it is proposed that our empathic experience involves a dynamic interplay between the two components with an explicit representation of another person’s affective state being a prerequisite, thereby making initial cognitive empathy necessary for empathic empathy to occur (Decety, & Jackson, 2004; Hillis, 2014).

Numerous studies have focused specifically on empathy in response to physical harm and neural networks responding to empathy for others in pain overlap substantially with those engaged during actual experience of pain (Marsh, 2018). Similar to physical pain, being socially rejected or excluded by others causes painful experience in humans and both share common neural circuits (Eisenberger et al., 2003). Moreover, observing others being excluded evokes lower need satisfactions and negative affect in the same way as when being excluded personally (Wesselmann, Bagg, & Williams, 2009). Empathy for physical pain represents a prototypical example of emotional contagion driven primarily by automatic bottom-up processes such as described in the perception-action mechanistic model (PAM) (Preston, & de Waal, 2002). In contrast, empathy for social pain requires a greater cognitive component, particularly mentalizing and theory of mind to integrate the complex social information (Bruneau, Pluta, & Saxe, 2012). This difference is also reflected in the underlying neural networks, with empathic responses to physical pain engaging regions involved in autonomic arousal and emotional reactivity such as the dorsal anterior cingulate (dACC) and the anterior insula (AI) (Lamm, Decety, & Singer, 2011; Yao et al., 2016), whereas empathy for social exclusion additionally engages regions involved in mentalizing and theory of mind such as dorsomedial, medial and ventromedial prefrontal cortex (DMPFC, MPFC and VMPFC) (Masten, Morelli, & Eisenberger, 2011).

Empathy for those in suffering leads to distressed feelings and a motivation for helping behavior (FeldmanHall, Dalgleish, Evans, & Mobbs, 2015). Indeed, the motivation for altruistic and prosocial behavior is an evolutionary outcome of empathy (Rumble, Van Lange, & Parks, 2010; de Waal, 2008). In line with this strong motivational component a previous study demonstrated that reciprocal altruism was associated with increased activity in the reward system, suggesting that it may be reinforcing (Rilling et al., 2002). However, in everyday life altruism is often exhibited in the absence of an expected reciprocity and sometimes even occurs at the cost of an individual’s self-interest (de Waal, 2008). Batson proposed the empathy-altruism hypothesis that empathic concern referring to other-oriented emotions produced altruistic motivational states with the goal of increasing the welfare of others rather than self (Batson, 2011). This costly altruism is predominantly exhibited towards genetically and socially close others but might also extend to strangers (Vekaria, Brethel-Haurwitz, Cardinale, Stoycos, & Marsh, 2017). Furthermore, costly altruistic behaviors and associated activity in reward-related regions are driven by other-oriented empathy rather than experienced personal distress (FeldmanHall, Dalgleish, & Mobbs, 2015). In social contexts altruistic punishment represents another form of costly altruistic behavior that aims both to help the victim and to enforce group norms and cooperation (West, Griffin, & Gardner, 2007; Boyd, Gintis, Bowles, & Richerson, 2003). Both forms of costly altruism have been associated with activity in core reward processing regions including the ventral striatum (De Quervain, Fischbacher, Treyer, & Schellhammer, 2004).

The hypothalamic neuropeptide oxytocin (OXT) has been found to influence a number of different aspects of social cognition and to promote the formation and maintenance of social bonds (Kendrick et al., 2017; Striepens, Kendrick, Maier, & Hurlemann, 2011; Macdonald, & Macdonald, 2010). Studies administering intranasal OXT have shown that it modulates core pain empathy regions, including the ACC and insula, as well as mentalizing regions and reward-related striato-frontal circuits although in a context-dependent manner (see Wigton et al., 2015; Bethlehem, van Honk, Auyeung, & Baron-Cohen, 2013). A number of genetic studies have also reported links between OXT-receptor polymorphisms and empathy in both Caucasian and Chinese populations (Rodrigues, Saslow, Garcia, John, & Keltner, 2009; Smith, Porges, Norman, Connelly, & Decety, 2014; Wu, Li, & Su, 2012). Intranasal OXT has been shown to particularly enhance emotional - rather than cognitive - empathy in both Caucasian and Chinese subjects (Geng et al., 2018; Hurlemann et al., 2010) and to concomitantly reduce amygdala reactivity (Geng et al., 2018; Becker et al., 2018). On the other hand, insula responses can be either enhanced (Riem et al., 2011; Striepens et al., 2012), or decreased in the context of pain empathy (Bos, Montoya, Hermans, Kevsers, & van Honk, 2015) and embarrassment (Becker et al., 2018).

In line with the emotional empathy-enhancing effects of intranasal OXT, a previous study has reported that it also increased altruistic behavior towards an ostracized individual (Riem et al., 2013). However, in this study exhibiting altruistic behavior did not come at any cost to the participants whereas in real-life situations it often does, and particularly in contexts where it might result in reduced financial gain or other aspects that conflict with self-interest (Camerer, & Fehr, 2006; Miller D, 2001). Although many prosocial effects of OXT have been reported it has also been found to promote self-serving lying and group-serving dishonesty (Sindermann, 2018, Shalvi & De Dreu, 2014). In addition, OXT’s detrimental effect on honesty was found in a competitive environment and driven by conformity with the behavior of peers (Aydogan, Jobst, D’Ardenne, Müller, & Kocher, 2017). On the other hand, intranasal OXT also tends to increase both altruistic help as well as costly altruistic (altruistic punishment) behaviors (Hu et al., 2016; Aydogan, Furtner, et al., 2017). Thus, although a number of studies have demonstrated the effects of OXT on empathy, altruistic and self-serving behaviors it is unclear what its functional role may be at both the behavioral and neural level when these motivations are competing in a social situation.

The current placebo-controlled double-blind fMRI study therefore aimed at determining the effects of intranasal OXT on competing behavioral tendencies between empathy-motivated altruism and self-interest. To this end a modified Cyberball paradigm was employed during which subjects initially observed a social exclusion situation and subsequently engaged in the game. The Cyberball game is a classic and widely used paradigm to induce social exclusion which leads to aversive and painful feelings by threatening an individual’s four fundamental needs including belonging, control, self-esteem and meaningful existence (Williams, 2009). The negative emotional responses to being socially excluded are found even when individuals know that they were excluded by a computer (Zadro, Williams, & Richardson, 2004). Brain imaging studies have demonstrated that the neural circuits that mediate exclusion-induced psychological pain resemble those responding to physical pain with common activations seen in the pain (e.g. anterior cingulate cortex, ACC) (Eisenberger et al., 2003) and somatosensory processing (e.g. secondary somatosensory cortex, dorsal posterior insula) systems (Kross, Berman, Mischel, Smith, & Wager, 2011). The Cyberball paradigm has also successfully been employed to induce social pain empathy by observing others being excluded in this game (Wesselmann, Bagg, & Williams, 2009) and subsequent prosocial behavior towards the excluded individual (Masten et al., 2011). Finally, previous studies have demonstrated a high sensitivity of this paradigm to capture effects of OXT on social behavior in both healthy subjects (Xu et al., 2017) and individuals with autism (Andari et al., 2010).

In the current adaptation of the Cyberball paradigm, participants were scanned while first observing 3 unknown individuals playing the game and where one player (victim) was gradually excluded by the two other players (excluders) so that they gained more money than the victim. Immediately after the observe session, participants were given an opportunity to play with the victim and one of the excluders as well as another new player. To create a situation of competing empathy and self-interest motivated behavioral tendencies participants were told that any player in this paradigm receiving a ball would receive an additional monetary reward (0.3 RMB/ball). In this case, the excluder player was manipulated to be the most attractive cooperator to maximize self-interest while playing with the victim of exclusion would be rather motivated by empathy-induced altruistic behavior.

Considering convergent evidence for OXT-enhanced emotional empathy, we hypothesized that it would increase empathy for the victim of exclusion during the first stage of the experiment. We additionally hypothesized that OXT-facilitated empathy would be accompanied by increased activation in (social) pain (dACC and anterior insula) and mentalizing networks (medial frontal, posterior parietal and temporal regions, i.e. posterior superior temporal sulcus (pSTS), posterior cingulate cortex (PCC) and precuneus (Eisenberger et al., 2003; Masten et al., 2011; Singer, 2006). In the second stage of the experiment when the subjects were actively engaged, we hypothesized that if OXT promotes costly altruistic behavior subjects should increase the proportion of their throws to the victim and correspondingly throw less to the excluder, whereas if it promotes increased self-serving behavior then this should result in a greater proportion of throws to the excluder player who should be more likely to reciprocate, thus increasing the financial gain. Given the engagement of striato-orbitofrontal reward processing circuits (Singer, 2006; Kringelbach, 2005; Spicer et al., 2007) in both monetary reward-anticipation (O’Doherty, Kringelbach, Rolls, Hornak, & Andrews, 2001) as well as altruistic behavior (FeldmanHall, Dalgleish, Evans, & Mobbs, 2015) we hypothesized that the OXT-induced behavioral preference for a player would be mirrored by increased activity in this circuit. Given that cultural orientation, i.e. horizontal independence modulated the effect of OXT following social exclusion (Xu et al., 2017), and higher trait altruism has been associated with stronger empathic brain responses (Haas et al., 2015) these traits were additionally assessed. Finally, in view of our previous study demonstrating long-term effects on memory for and preference for replaying with specific players (Xu et al., 2017) we also investigated these same factors one week after the social exclusion experiment.

## Materials and Methods

### Participants

82 healthy Chinese male university students (right-handed, age = 18-27 years, Mean ± SE = 21.36 ± 0.24 years) participated in the present study and were randomly assigned to receive either placebo (PLC) (n = 41, age = 18-26 years, Mean ± SE = 21.68 ± 0.34 years) or OXT nasal spray (n = 41, age = 18-27 years, Mean ± SE = 21.05 ± 0.34 years; PLC vs. OXT, *t* = 1.31, *p* = 0.195) in a double-blind between-subject pharmacological fMRI experiment. Subjects reported being free from current or a history of psychiatric or neurological disorders and did not use any medication in the four weeks before the experiment and were asked to abstain from caffeine and alcohol in the 24 hours before the experiment.

Study procedures were approved by the local ethics committee at the UESTC and adhered to the latest revision of the Declaration of Helsinki. Each participant provided written informed consent before the experiment and received monetary compensation for participation. Study protocols were pre-registered at clinical trials.gov (https://www.clinicaltrials.gov/ct2/show/NCT03122067, Trial ID: NCT03122067).

### Procedure

To control for potential confounding effects of pre-treatment differences in affective state and empathy-related domains, validated Chinese versions of established questionnaires to assess mood (Positive and Negative Affect Schedule, PANAS) (Watson, & Clark, 1988), anxiety (State-Trait Anxiety Inventory, STAI), (Barnes, Harp, & Jung, 2002), (Liebowitz Social Anxiety Scale, LSAS) (Heimberg et al., 1999), depression (Beck Depression Inventory, BDI) (Beck, Steer, & Brown, 1996), trait autism (Adult Autism Spectrum Quotient, ASQ) (Baron-Cohen, Wheelwright, Skinner, Martin, & Clubley, 2001), trait empathy (Interpersonal Reactivity Index-C, IRI-C) (Siu, & Shek, 2005), and early life stress (Childhood Trauma Questionnaire, CTQ) (Bernstein, & Fink, 1998) were administered. In line with our previous study on the effects of OXT on social exclusion (Xu et al., 2017) we explored associations with the ‘horizontal independence’ (HI) scale of the Individualism and Collectivism Scale (ICS) (Singelis, Triandis, Bhawuk, & Gelfand, 1995). In the context of previously reported associations between trait altruism and empathic brain activity (Haas et al., 2015), as well as a potential contribution of altruism-driven attempts to compensate the victim after observing them being excluded we additionally explored associations between the effects of OXT and the level of pre-treatment altruistic prosocial behavior as assessed by the altruistic subscale of the Prosocial Tendency Measure (PTM). This altruistic subscale in the PTM specifically assesses concerns about individuals in need of help that incurs a cost to the helper (Carlo, & Randall, 2002) and thus appears of particular relevance with respect to the present paradigm. Mood (PANAS) was additionally assessed after the experiment to control for unspecific effect of treatment on these domains.

After participants completed the questionnaires portrait photos with neutral facial expressions were taken from each subject and served as stimuli depicting them during the subsequent Cyberball game. Finally, subjects were asked to rate portrait photos with neutral facial expressions of their subsequent fellow-players in the experiment with respect to likeability, trustworthiness and valence (for details see paradigm description). Subjects next self-administered 24 IU OXT (Oxytocin-spray, Sichuan Meike Pharmaceutical Co., Ltd; 3 puffs of 4IU per nostril with 30s between each puff) or placebo (PLC – identical sprays with the same ingredients other than OXT – i.e. sodium chloride and glycerol) 45 minutes before the start of the experimental paradigm. Administration adhered to a standardized protocol for the intranasal administration of OXT (Guastella et al., 2013).

### Modified Cyberball paradigm

Participants were told they would first observe and then participate in a ball-tossing game (‘Cyberball’) with 4 other participants online while they were in the fMRI scanner and that the other 4 players were sitting in separate compartments in a nearby behavioral testing room to avoid direct personal interaction during the entire experiment. However, in fact the 4 players were fictitious and preprogrammed in the experimental paradigm. The revised Cyberball task in this study included two sessions acquired during separate fMRI runs. The first session (OBSERVE condition) aimed to prime the subject’s attitude towards each of the players observed (victim or excluder, see Fig. 1a) in order to influence their subsequent behavior when they participated in the game during the second session (PLAY condition, see Fig. 1b). During the OBSERVE run, subjects simply observed 3 individuals playing the game and were told that each player in the game would receive 0.3RMB reward for every ball thrown to them. Subjects were explicitly instructed to observe the game and to consider what each player might be thinking or feeling during it (Masten et al., 2011). The first run (OBSERVE) contained 10 blocks with each block lasting for 30 seconds, starting with 4 ‘fair’ blocks during which the 3 players threw the ball equally often to each other, followed by two ‘transition’ blocks during which one of the players was gradually excluded and 4 ‘exclusion’ blocks during which one player (victim) was totally excluded by the other two players (excluder 1 and excluder 2). After each round the subject was shown how much money each of the 3 players had gained (displayed for 12 s). In this way the subject observing the game was expected to detect that one of the players (victim) was eventually excluded by the two other players (excluders) and that the result of this was the two excluders had learned that by throwing only to each other they ended up gaining more money at the expense of the victim who gained less. The second run (PLAY) involved the subject and 3 other players, including the victim, one excluder (1 or 2) from the first run and a new player and comprised 6 rounds with a fixed number of 24 ball throws per round. All virtual players were programmed to throw the balls equally to the other 3 players. Subjects were told that during the first four rounds they would be informed how much each player would have gained (0.3RMB per ball received – amount gained per player was displayed for 12 s) but that they would only actually receive the monetary reward from the final 2 rounds. This was done to allow us to assess whether having a strong motivational component of monetary rewards in every round might have biased behavior towards greater self-interest behavior and reduce altruistic behavior towards the victim from the first game. To confirm whether the pattern of throwing to the other players was different when a monetary incentive was included, we initially explored potential interaction effects between treatment and the monetary condition (i.e. first 4 vs. last two blocks). In line with our expectation no main effect of monetary condition or interaction effects with treatment were found (for details see supplementary information) and consequently this factor was discarded from all further analyses.

**Figure 1.**
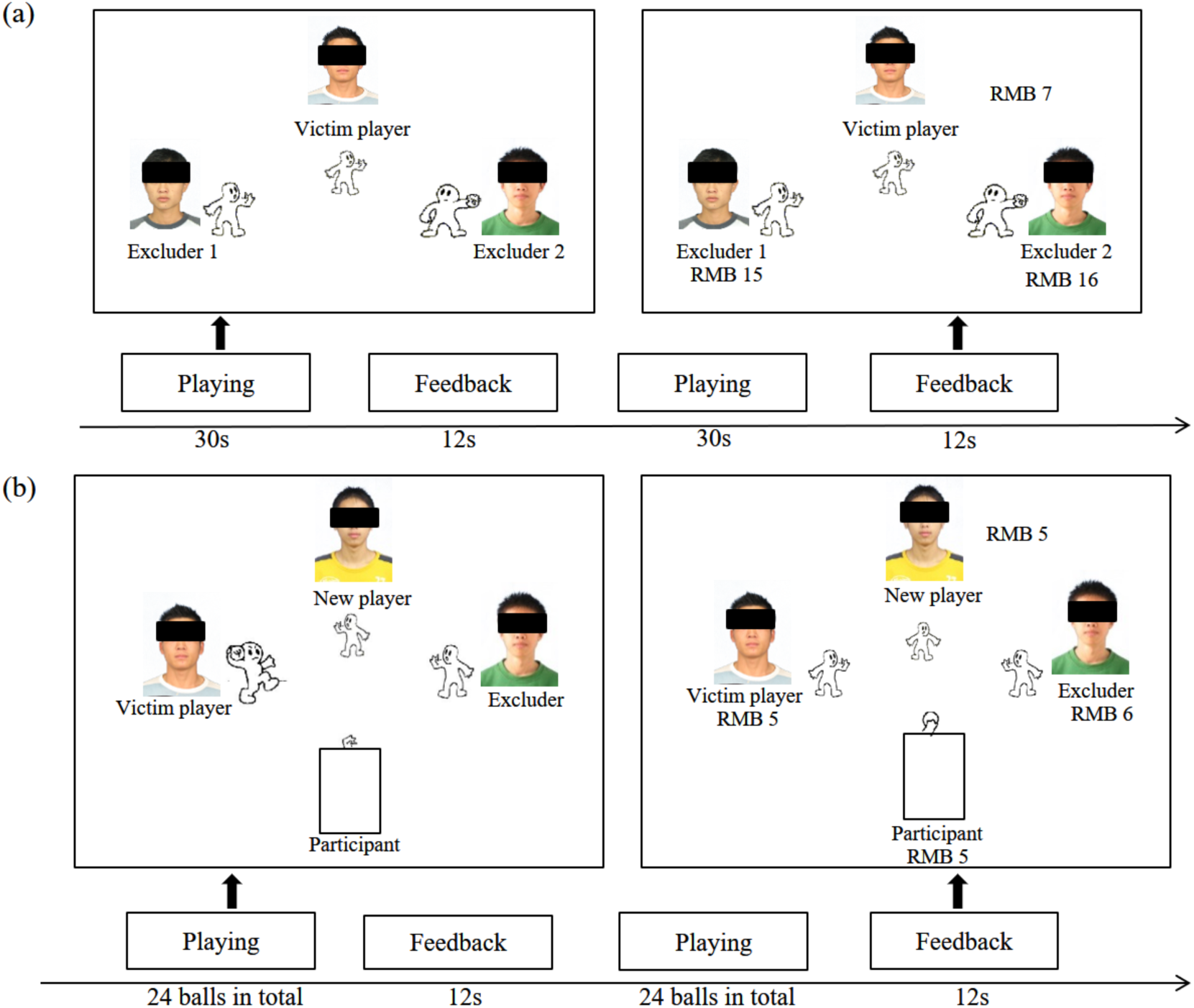
The Cyberball game paradigm employed. (a) Subjects first completed an “Observe” session which included a total of 10 blocks (4 blocks where players threw equally to each other followed by 2 blocks where two players (excluders) started to throw more often to each other and finally 4 blocks where the excluders threw exclusively to each other and did not throw to the other player (victim) at all). Blocks lasted for 30s during a 12s period between each block the subjects viewed how much money each of the players had won. Subjects were instructed during their observation of the game to consider what the individuals playing it were thinking and feeling. (b) Subjects next completed a “Play” session where they played the game together with the victim and one of the excluders and a novel player. This session was for 6 blocks with 24 ball throws in each block (After each block the subject could see how much money they and the other players had won and a 2 s maximum was allowed to throw the ball to avoid a monetary deduction).

### Manipulation check and potential confounders

As a manipulation check subjects were asked to rate the other players in terms of their likeability, trustworthiness and valence using a visual analogue scale (VAS, 0-100) before treatment administration and following the OBSERVE and PLAY sessions respectively. To further evaluate the manipulation subjects were asked to report if any specific events had happened during the OBSERVE session (e.g. “Did all three players treat each other fairly?”, “Was there anyone who was treated unfairly?”). They were also asked to rate how empathic they felt towards the excluded player (score between 0-100). As additional control variables the PANAS questionnaire was administered again to test whether OXT had altered participants’ mood (PANAS) after the paradigm.

### Long term effects of OXT – follow-up after one week

Subjects were asked to return to the laboratory to complete follow-up assessments one week after the Cyberball game. The assessment included ratings of the other players with respect to likeability, trustworthiness and valence. Moreover, a surprise memory test was employed to examine if the previous Cyberball game had an effect on social recognition memory and whether this was influenced by OXT. Finally, subjects were asked if they were willing to play with the previous players again and rated how much they would like to do so (1-9 scale).

### Behavioral data analyses

Statistical analyses for the questionnaires and behavioral ratings were performed using SPSS 18.0 software (SPSS Inc., Chicago, Illinois, USA). Post-hoc analyses of interaction effects were performed employing Bonferroni correction for multiple comparisons. Associations between traits, behavior and neural indices were examined using Pearson correlation and differences in the correlations between the two groups were further examined using Fisher’s Z test with Bonferroni correction.

### Image acquisition

Imaging data were collected using a 3T GE Discovery MR750 system (General Electric, Milwaukee, WI, USA) using the following sequence parameters: TR = 2000 ms; TE = 30 ms; flip angle = 90°; number of slices = 43; slice thickness = 3.2 mm; FOV = 220 × 220 mm2; matrix = 64 × 64; slice orientation = axial. High-resolution whole-brain T1-weighted images were additionally acquired using a spoiled gradient echo pulse sequence to improve normalization of the functional data (TR = 6 ms; TE = 2 ms; flip angle = 12°; number of slices = 156; slice thickness = 1 mm; FOV = 256 × 256 mm2; matrix = 256 × 256).

### fMRI data analysis

Statistical Parametric Mapping as implemented in SPM12 (http://www.fil.ion.ucl.ac.uk/spm/) was used to preprocess and analyze the neuroimaging data. The first 6 volumes of each functional neuroimaging time-series were removed to allow for T1 equilibration. Preprocessing included slice timing, image realignment to correct for head motion, normalization into the Montreal Neurological Institute (MNI) space resampled at 3×3×3 mm voxel size, and spatially smoothed using an 8 mm FWHM Gaussian kernel. Generalized linear models (GLM) were built to investigate the BOLD signal changes. A 128-second high-pass filter was applied to further control for low-frequency noise artifacts.

The first-level design matrix for the OBSERVE run was modelled using a blocked-design matrix including the first 4 blocks as the inclusion condition and the last 4 blocks as the exclusion condition. The monetary reward feedback and the middle two blocks with the transfer between inclusion and exclusion were additionally modelled and the six head motion parameters included.

The first-level design matrix for the PLAY run was modelled using an event related design matrix to specifically examine the throws made by the participant to the other individual players. Ball-tosses towards the excluder, victim and new player were implemented as experimental conditions and modelled as separate events. To specifically model the expectation phase for a reciprocal action, independent from the decision phase, the time between the other player receiving the ball form the participant and the throw of that player was modelled as experimental event. Monetary feedback, rating periods and head motion parameters were additionally included in the matrix. Due to technical issues during the fMRI assessment and excessive motion (> 3mm), data from 8 participants had to be excluded (OXT = 4, PLC = 4), leading to a final sample size of OXT =37 and PLC = 37 for the imaging analysis. Effects of OXT during the OBSERVE and PLAY sessions were assessed by employing independent t-tests. In line with the main aim of the study the contrast [exclusion > inclusion] was used for the OBSERVE condition, and for the PLAY condition player-specific contrasts were examined [excluder, victim, new player]. The threshold p-value level was set at <0.05 cluster-level with family wise error (FWE) correction for multiple comparisons and an initial cluster forming threshold at the voxel-level of *p*<0.001, uncorrected (in line with recommendations by Eklund et al., 2016; Slotnick; 2017)

## Results

### Potential confounders

No significant differences between subjects in the PLC and OXT groups were found in pre-treatment affective state and empathy-related domains (Independent t-tests, Table 1). Importantly, mood as assessed by the PANAS did not differ after the experiment, arguing against unspecific confounding effects of treatment on mood.

**Table 1.** Questionnaire scores in PLC and OXT group before and after treatment.

| Measurements | Placebo | Oxytocin | *t*-value | *p*-value |
| --- | --- | --- | --- | --- |
| **Before treatment** |  |  |  |  |
| Positive and Negative Affect Schedule (PANAS) |  |  |  |  |
| Positive | 29.15±1.10 | 28.93±0.79 | 0.16 | 0.87 |
| Negative | 15.20±0.79 | 14.29±0.75 | 0.83 | 0.41 |
| State-Trait Anxiety Inventory (STAI) |  |  |  |  |
| State | 37.68±1.44 | 36.93±1.32 | 0.39 | 0.70 |
| Trait | 37.63±1.20 | 39.56±1.22 | -1.13 | 0.26 |
| Liebowitz Social Anxiety Scale (LSAS) |  |  |  |  |
| Avoid | 19.85±1.40 | 21.56±1.31 | -0.89 | 0.38 |
| Fear | 22.56±1.53 | 20.2±1.34 | 1.16 | 0.25 |
| Beck Depression Inventory (BDI-II) | 5.41±1.32 | 6.10±1.09 | -0.40 | 0.69 |
| Adult Autism Spectrum Quotient (ASQ) | 20.51±0.70 | 18.76±0.80 | 1.64 | 0.10 |
| Interpersonal Reactivity Index (IRI) | 39.76±0.99 | 40.29±1.03 | -0.38 | 0.71 |
| Childhood Trauma Questionnaire (CTQ) | 56.20±0.80 | 56.00±0.91 | 0.16 | 0.87 |
| Individualism and Collectivism Scale (ICS) | 5.31±0.11 | 5.36±0.12 | -0.31 | 0.76 |
| Prosocial Tendency Measure (PTM)-altruism | 15.93±0.43 | 16.76±0.40 | -1.41 | 0.16 |
| **After treatment** |  |  |  |  |
| PANAS-Positive | 25.68±1.48 | 24.46±1.19 | 0.64 | 0.52 |
| PANAS-Negative | 12.85±0.57 | 13.60±0.86 | -0.73 | 0.47 |

### Effects of perceived social exclusion and oxytocin on likeability, trustworthiness and valence ratings

A 2 (Treatment: OXT/PLC) × 3 (Player: Excluder/Victim/New) mixed ANOVA with subjects’ likeability, trustworthiness and valence rating scores as dependent variables revealed no significant differences in these three dimensions between the PLC and OXT groups before the experiment (Fig. 2a). However, after the OBSERVE session during which subjects watched the victim player being excluded, their ratings of the victim player for the three dimensions were significantly higher than for the excluders (Fig. 2b), confirming successful experimental manipulation. Moreover, the decreased ratings for the excluder players remained stable after subjects played with them during the PLAY run and were even maintained one week later (Figs. 2c, 2d).

**Figure 2.**
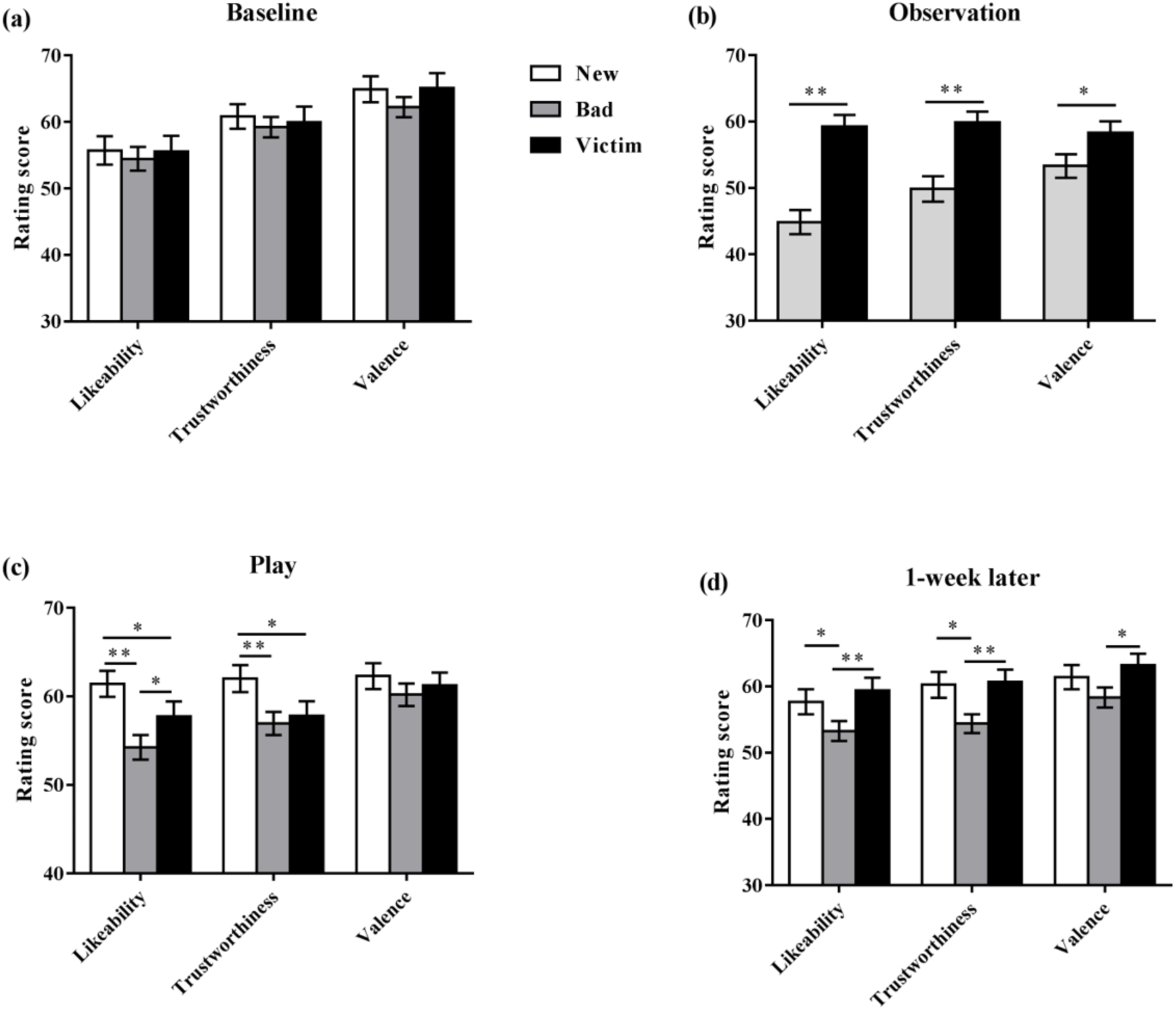
Rating scores of likeability, trustworthiness and valence in combined oxytocin and placebo-treated groups for (a) before treatment, (b) after treatment during the OBSERVE session, (c) after PLAY session, (d) 1-week after treatment and observing and playing the Cyberball game. \**p*<0.05, \*\**p*<0.005.

### Manipulation check and empathic responses

At the end of the experiment all subjects reported that they realized one player was excluded during the OBSERVE session further confirming successful manipulation. To examine the effects of OXT on subjects’ empathy scores for the victim player, an independent two sample t-test was performed but revealed no significant differences between the treatment groups (PLC = 63.62, OXT = 58.97, *t* = 0.98, *p* = 0.33, Cohen’s *d* = 0.224).

### Behavior during the PLAY phase of the paradigm

For the PLAY condition, the percentage of ball tosses to each of the other three players served as the dependent variable and was subjected to a 2 (Treatment: PLC/OXT) × 3 (Players: Excluder/Victim/New) mixed ANOVA. Results revealed a main effect of player (*F*_*2,160*_ =7.05, *p* = 0.001, *η^2^_p_*=0.081), and a marginal significant interaction effect between treatment and player (*F*_*2,160*_ = 2.74, *p* = 0.068, *η^2^_p_* =0.033). Post-hoc Bonferroni-corrected paired comparisons showed that subjects in both groups threw more balls to the victim than to the excluder players (victim = 36.7%, excluder = 30.28%, *p* = 0.002, Cohen’s *d* = 0.690). Further exploratory analysis of the interaction effect demonstrated that the OXT group threw significantly more balls to the excluder player relative to the PLC group (to excluder player: PLC = 27.97%, OXT = 32.60%, *t* = −2.46, *p* = 0.016, Cohen’s *d* = 0.543, Fig. 3).

**Figure 3.**
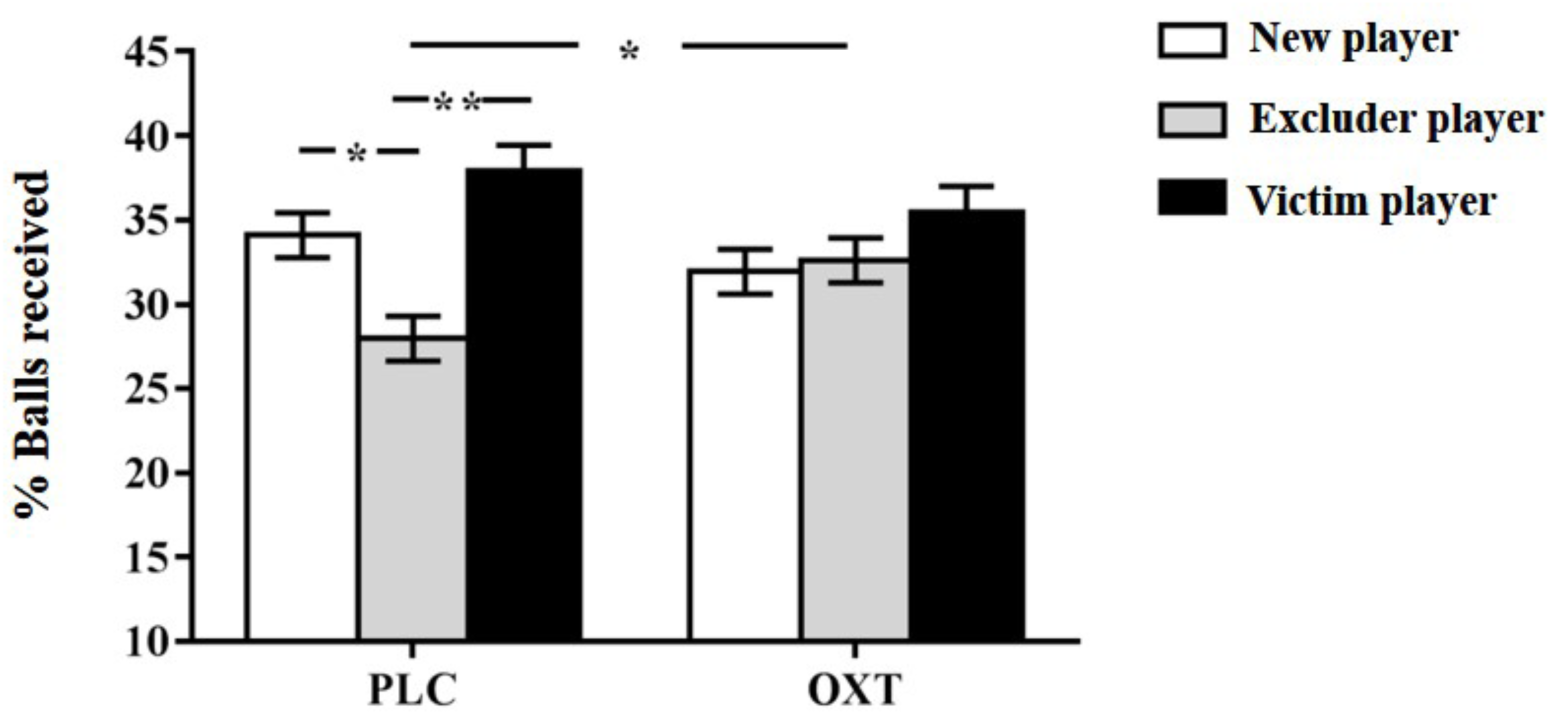
Percentage (mean ± sem) of balls that participants threw to the excluder, new, and victim players respectively. \**p*<0.05, \*\**p*<0.005.

### Neural activation changes during the Cyberball task

The task-related network engaged during the observation of social exclusion and effects of OXT on the underlying neural activity [contrast, exclusion > inclusion] were firstly investigated during the OBSERVE session. Analysis of neural changes during the OBSERVE session in the combined PLC and OXT groups revealed significant increased activity in the posterior cingulate cortex (PCC), left midcingulate cortex (lMCC), precuneus, left inferior parietal lobule (lIPL) and right posterior superior temporal sulcus (rpSTS) when subjects observed the victim being excluded (Fig. 4, table 2). In line with the lack of OXT effects on the post OBSERVE behavioral ratings, no significant neural differences were observed between the treatment groups. Thus, any group differences during the subsequent PLAY session were unlikely to be driven by effects of OXT during the preceding encoding phase of the social interaction.

**Table 2.** Brain regions that positively activated under observed exclusion > inclusion. *p*<0.05 (FWE corrected, cluster level)

| Regions | Cluster k | x | y | z | t |
| --- | --- | --- | --- | --- | --- |
| **Posterior cingulate cortex** | 2100 | -3 | -84 | 24 | 7.00 |
| Midcingulate cortex, L |  | 0 | -36 | 39 | 6.70 |
| Precuneus, L |  | -9 | -51 | 3 | 6.57 |
| Inferior parietal lobule, L | 169 | -42 | -75 | 39 | 6.35 |
| Superior temporal sulcus, R | 257 | 60 | 0 | 3 | 5.82 |
|  |  | 60 | -18 | 9 | 3.60 |
| Rolandic operculum, R |  | 48 | -15 | 15 | 4.23 |

**Figure 4.**
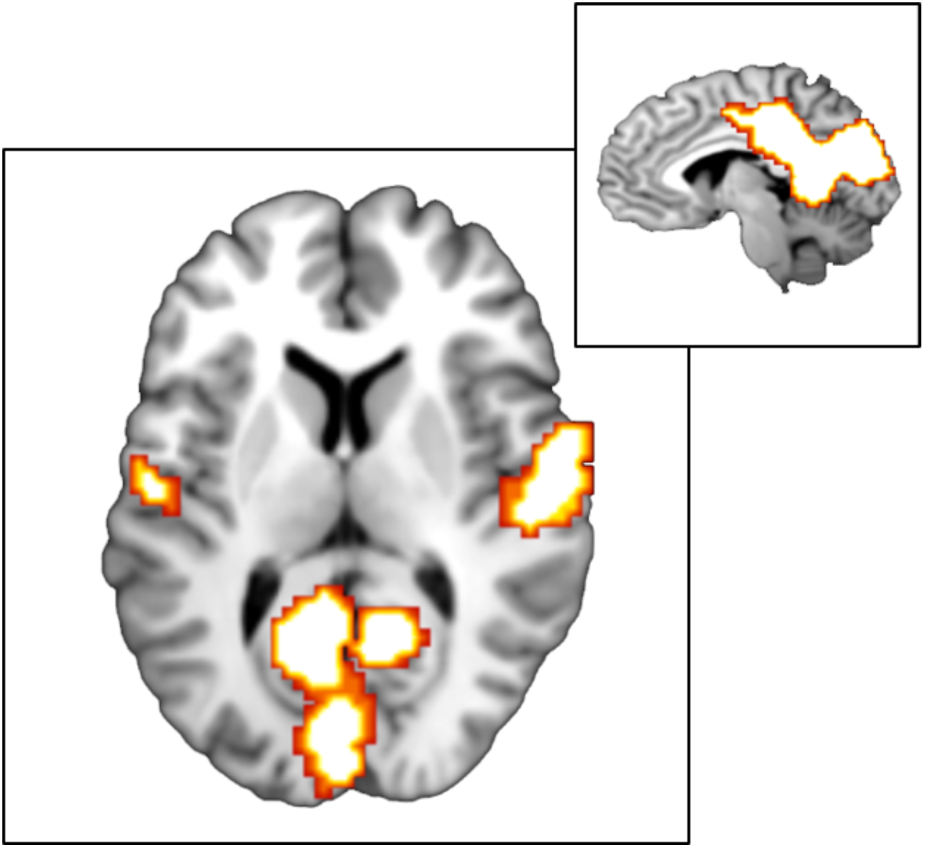
Neural responses to observed exclusion > inclusion in the entire sample (i.e. oxytocin and placebo treated groups combined – *p* < 0.05; FWE corrected, cluster level).

For the PLAY session, a 2 (Treatment: PLC/OXT) × 3 (Player: Excluder/Victim/New) mixed ANOVA using the flexible factorial model in SPM was performed on the whole brain level but no significant results were found (FWE-corrected *p* < 0.05). Based on the behavioral results indicating that OXT administration specifically increased the proportion of throws made to the excluder player, an independent t-test was performed to examine neural activity differences between the treatment groups during this condition. Results indicated significantly stronger left medial orbitofrontal cortex (mOFC) activity (*k* = 106, *t* = 4.40, *x/y/z*: -6/24/-6, *p* = 0.047, Fig. 5) when playing with excluder players in the OXT group compared to the PLC group (FWE-corrected *p* < 0.05). In line with the behavioral findings exploratory t-tests revealed no significant treatment effects on neural activity when playing with the victim or the new player (both *p*s > 0.348).

**Figure 5.**
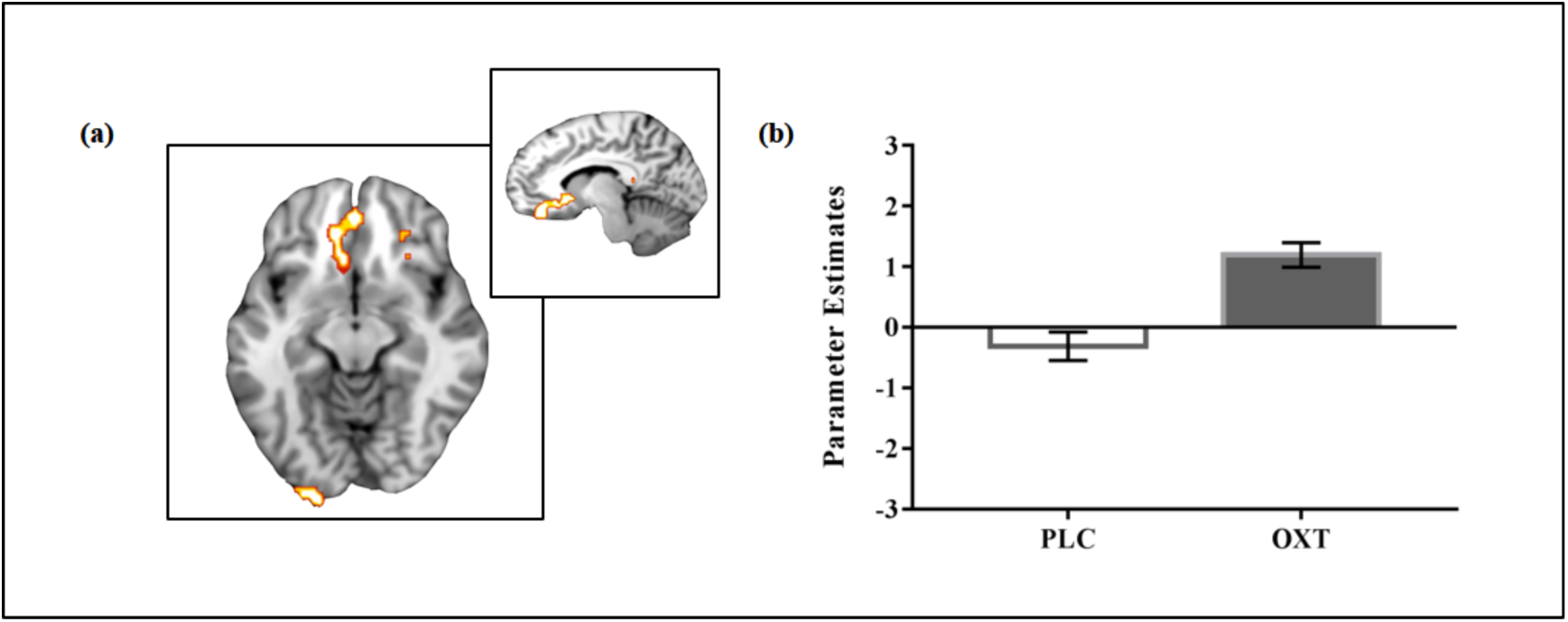
(a) Neural response to excluder player under oxytocin (OXT) > placebo (PLC). *p* < 0.05 (FWE corrected, cluster level), (b) Corresponding beta parameters of the left mOFC in the PLC and OXT group.

### Associations between trait altruism behavior ratings and neural activity

A potential modulatory influence of pre-treatment variations in trait altruism (PTM scores) on behavioral and neural responses was investigated using a correlation analysis. No significant associations between the proportion of balls thrown to the three different players and PTM scores were found in either the PLC (Excluder – r = -0.172, *p* = 0.283; Victim – r = 0.091, *p* = 0.57; New – r = 0.075, *p* = 0.64) or OXT (Excluder – r = -0.131, *p* = 0.413; Victim – r = 0.227, *p* = 0.154; New – r = -0.114, *p* = 0.478) groups, or differences between the groups (all *p*s>0.41). Correlations between left mOFC activity and PTM scores during play with the excluder were also examined in the PLC and OXT groups. Results revealed that PTM scores were not associated with mOFC activity following PLC, but that OXT treatment established a significant negative association between PTM trait altruism and activity in this region (PLC: r = 0.294, *p* = 0.094; OXT: r = -0.329, *p* = 0.047; significant different correlations confirmed by Fisher’s z = 2.59, *p* = 0.040, Cohen’s *q* = 0.28, Bonferroni corrected, Fig. 6). This shows that OXT particularly increased left mOFC activity in individuals with lower trait altruism. Horizontal Independence was not associated with either behavior or mOFC activity.

**Figure 6.**
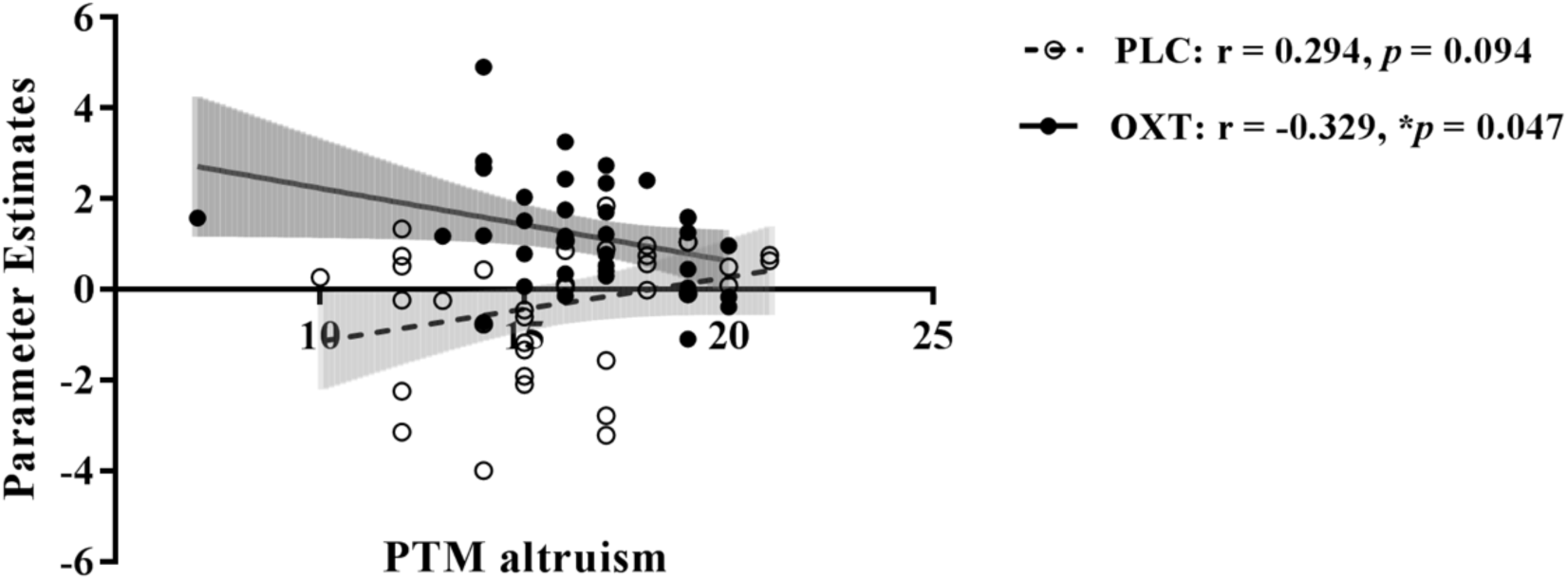
Correlations between pre-treatment trait altruism score (PTM) and neural activity when interacting with the excluder player in the placebo (PLC) and oxytocin (OXT) treatment groups. Best-fit line with 95% confidence bands, \**p* < 0.05

### Follow-up assessment after one week

Accuracy on memory for the faces of the different players was assessed in the two treatment groups during a surprise recognition memory test one week after the initial Cyberball games, however both groups achieved a very high accuracy (all OXT treated subjects scored 100% accuracy as did 38/41 of the PLC treated ones) and so no meaningful statistical comparison could be made. A 2 (Treatment: PLC/OXT) × 3 (Player: Excluder/Victim/New) mixed ANOVA using the willingness to replay rating scores as a dependent variable revealed a main effect of player (*F*_*2,160*_ = 10.46, *p* < 0.001, *η^2^_p_* = 0.116), with subjects reporting a stronger preference to play with the victim and new players again as compared to the excluder players (excluder = 3.36±0.27 vs. victim = 4.89±0.32, *p* = 0.002, Cohen’s *d* = 0.562; excluder vs. new = 4.93±0.29, *p* < 0.001, Cohen’s *d* = 0.611). A treatment × player interaction (*F*_*2,160*_ = 3.55, *p* = 0.031, *η^2^_p_* = 0.042,) showed additionally that whereas subjects in the PLC group were significantly less willing to play again with excluders compared to the victim or new players (excluder vs. victim, excluder vs. new, both post hoc *p*s < .005) this was not the case in the OXT group (both *p*s > 0.097).

## Discussion

The current study aimed firstly to establish whether OXT treatment enhanced empathic behavior and neural responses towards observing that a person (victim) is being socially excluded, and secondly whether it promoted altruistic or self-serving behaviors and associated neural responses under circumstances when these two behaviors are in competition with one another. Overall, on the behavioral level following observation of a modified Cyberball game both groups showed strong empathic responses towards the victim players and also greater likeability and trustworthiness ratings for them compared with excluder players, although OXT did not potentiate this. On the neural level, observation of social exclusion was accompanied by increased activity in the mentalizing network, including core regions such as PCC, pSTS, IPL and precuneus, however, in line with the lack of behavioral OXT effects, neural activity patterns were not influenced by OXT treatment. During the subsequent play phase of the paradigm subjects in both treatment groups threw more balls to the victim player but this effect was not enhanced by OXT, suggesting that it did not promote greater altruistic behavior. However, the OXT group threw significantly more balls to the excluder player suggesting that OXT promoted self-serving decisions since playing with the excluder should lead to a higher monetary pay-off. In line with our hypothesis, increased self-serving behavior following OXT was associated with stronger activation in the mOFC reward system when subjects were playing with the excluder player. Furthermore, OXT established a negative relationship between mOFC activity and trait altruism, an association that was absent following PLC. One week after the Cyberball game while the PLC group expressed a greater preference to play again with the victim and novel players compared to the excluder, in the OXT group there was no such difference indicating that they maintained their greater interest in playing with individuals who might potentially help them gain larger rewards. Thus overall, our results demonstrate that when altruistic and self-serving motivations are in competition OXT rather than promoting altruism actually enhances selfish decision making.

Our hypothesis that OXT would enhance empathic responses towards the victims of exclusion in the Cyberball game was not supported at either the behavioral or the neural level. Previous research combined OXT treatment with the multifaceted empathy task to demonstrate that it enhanced emotional but not cognitive empathy towards individuals in both positive and negative valence contexts (Hurlemann et al., 2010; Geng et al., 2018), an effect that was associated with suppressed amygdala responses (Geng et al., 2018). Oxytocin also increased empathic embarrassment in male and female subjects and this was associated with decreased amygdala and insula cortex responses, but with no effect on mentalizing networks (Becker et al., 2018). Empathic embarrassment can be considered as an example of social pain and another study has also reported that OXT decreased insula responses to viewing people in pain (Bos et al., 2015). In the current study in both the PLC and OXT groups there was evidence at the whole brain level for increased activation in core mentalizing regions (for convergent findings see also (Masten et al., 2011)) but no responses in the pain network (notably the insula).

In general, empathic ratings by subjects for the victim were not that high and since altered activation was only observed in the mentalizing network, which we have shown in the context of empathic embarrassment is not influenced by OXT (Becker et al., 2018), it seems possible that the empathy experienced in the current context was more cognitive than emotional. Indeed, given that the participants could not directly observe the other players and static neutral faces were used, the social pain had to be inferred by mentalizing how the victim may emotionally experience exclusion from the game. Together with the fact that mentalizing system is a core component of the cognitive empathy system (Shamay-Tsoory, 2011), this may explain why OXT failed to have any impact on empathic ratings or associated likeability and trustworthiness ratings for the victim in the current study since it has a greater influence on emotional rather than cognitive empathy.

Our original hypothesis that in a situation where altruistic and self-serving motivations were in competition OXT would enhance altruistic responses was also not supported in the current experiment. Overall, participants in both groups threw more balls to the victim player indicative of an altruistic response and validating the experimental manipulation, however OXT had no effect on this. Similarly, both groups rated the likeability and trustworthiness of excluder players lower than that of victims after observing the game and also one week later after playing it, but again this was not influenced by OXT. This general finding is in agreement with previous studies which also found that subjects exhibit greater prosocial behavior towards individuals who have been observed to be socially excluded (Masten et al., 2011; Van Der Meulen, Van IJzendoorn, & Crone, 2016). On the other hand, the OXT group threw a significantly greater proportion of balls to the excluder player indicative of an enhanced self-serving motivation since the excluder player would have been perceived as being more likely to reciprocate and potentially result in greater financial gain for the participant.

While a number of previous studies have demonstrated that OXT can facilitate altruistic behaviors in terms of cooperation, generosity, trait-judgements and valuing other’s possessions, these have mainly involveed contexts where personal costs to individuals were absent or low (Declerck, Boone, & Kiyonari, 2010; Andari et al., 2010; Riem et al., 2011; Zhao et al., 2016; 2017). The finding in the current study that under circumstances where there is perceived to be a potential cost of altruism in terms of reduced personal gain argues for the primary function of OXT as enhancing the motivation for resource acquisition. Where individuals do exhibit costly altruistic behavior, this is paralleled by increased empathic concern and altered activation in the ventral tegmental area, caudate and subgenual anterior cingulate which are important for promoting social attachment and caregiving (FeldmanHall, Dalgleish, Evans, & Mobbs, 2015). Although OXT has been show to modulate neural processing in these regions in social and non-social contexts (e.g. Scheele et al., 2013; Mickey et al., 2016; Zhao et al., 2019) it did not affect activity in this circuitry during the present paradigm, further indicating its lack of effect on promoting altruism in the current context.

Previous studies have reported that OXT can promote lying for the benefit of in-group members, including participants themselves, although not lying purely for self-gain (Shalvi, & De Dreu, 2014) and that it reduced honesty for personal gain only in a competitive environment (Aydogan et al., 2017). However, OXT can also in some circumstances promote pure self-serving lying to increase personal gain in men when there is no risk of discovery (Sindermann et al., 2018) and can increase acceptance of self-benefit moral dilemmas, but importantly not acceptance of other types of moral dilemma (Scheele et al., 2014). Interestingly, OXT effects on self-serving lying for financial gain are modulated by OXT receptor genotype (Sindermann et al., 2018) and thus it is possible that the effects of OXT on altruistic compared with self-benefit behaviors when they are in conflict might be to some extent genotype dependent. Taken together, and in line with our current results, accumulating evidence therefore suggests that OXT can promote personal self-interest in some contexts.

On the neural level the present study revealed that increased self-serving behavior following OXT was neurally underpinned by regional- and player-specific increased mOFC activation when participants interacted with excluder players. The mOFC is involved in monitoring associations between previous stimuli with reward and tracking response-outcome probabilities during changing reward contingencies (Elliott, Dolan, & Frith, 2000; Kringelbach & Rolls, 2004). Moreover, the mOFC codes the value of different behavioral options (Padoa-Schioppa, & Assad, 2006), including the value of expected monetary gains (Breiter, Aharon, Kaheman, Dale, & Shizgal, 2001), and activity in this region increases with monetary reward magnitude (O’Doherty, Kringelbach, Rolls, Hornak, & Andrews, 2001). Thus, in the current context increased mOFC responses may reflect an enhanced value of the expected greater monetary reward when cooperating with the excluder player in subjects in the OXT group.

Additionally, OXT treatment produced a negative association between PTM trait altruism and mOFC activation that was absent during PLC treatment. This suggests that at the neural level OXT particularly increased the value of the potential monetary gain in subjects with low baseline altruistic tendencies. Possibly individuals with high trait altruism might be less likely to experience a greater anticipation of gaining a greater monetary reward by playing with the excluder under OXT, as a result of greater feelings of guilt evoked by having to exclude the other players, and notably the victim.

While both behavioral and neural effects of OXT observed in the current paradigm do indicate a shift towards a self-serving rather than altruistic motivation it is notable that the pattern of altered bias is quite subtle. Under OXT, participants do not actually play more with the excluders than with either the victim or the novel player and effectively exhibit an egalitarian playing pattern which is therefore unlikely to create any feelings of exclusion in any of the other players. This is in contrast to participants in the PLC group who show a clear pattern of excluding the previous excluder by comparison with both the victim and novel players. As such the effect of OXT could be viewed as promoting self-benefit behavior but only if it does not damage others and therefore possibly generate significant feelings of guilt. Indeed, this is in accordance with the findings that OXT increases lying for self-gain when individuals know that there is no chance their lies will be discovered or that this comes at the cost of reducing the financial gain of others (Sinderman et al., 2018). Alternatively, it might be argued that subjects in the PLC group are exhibiting altruistic punishment towards the excluders and OXT is reducing the desire to inflict such punishment. However, altruistic punishment is strongly associated with altered amygdala function (Scheele et al., 2012) and there was no evidence for differential amygdala responses in the PLC and OXT groups. While a previous study has reported that OXT can promote altruistic punishment of defectors, and feelings of anger and disappointment towards them, in an economic game context, it also increases co-operation with them thereby increasing self-gain (Prisoner’s dilemma game: Aydogan et al., 2017). Thus, on balance, it is likely that in the current context OXT primarily biases individuals towards an optimal self-gain strategy, although without simultaneously doing so by overtly damaging others emotionally. Clearly, to further establish this it would be necessary to investigate the effects of OXT under circumstances where increasing self-gain would also significantly damage others emotionally.

There are several limitations in the present study which should be noted. Firstly, the victim observed being excluded in the Cyberball paradigm in the current study was a stranger to the participant and therefore it is possible that if they had been a partner, relative or other in-group member then OXT may have had the opposite effect by enhancing empathic and altruistic behaviors rather than self-serving ones. Further studies would be required to establish this. Secondly, and relatedly, only male participants were included in the current study and previous studies reporting OXT-induced enhancement of self-serving behaviors have also only included male subjects. A number of studies have reported opposite neural and behavioral effects of OXT in males and females (Gao et al., 2016; Luo et al., 2017) and one study found that while it enhanced acceptance of self-benefit moral dilemmas in men it had the opposite effect in women (Scheele et al., 2014). Thus, it remains possible that in a similar competing motivation situation OXT may have facilitated altruistic rather than self-serving behavior in women.

In summary, our current findings demonstrate that under circumstances where self-serving and altruistic behaviors are in competition OXT promotes an increased tendency in men towards self-benefit behavior, and this is associated with increased activation in the mOFC indicative of greater reward anticipation. Furthermore, the effects of OXT on mOFC are strongest in individuals with lower trait altruism. Thus, contrary to expectation, OXT did not promote altruistic behavior although equally it did not reduce it to the point where it might potentially have produced negative feelings in others. It seems likely therefore that in men OXT tends to bias individuals towards acquisition of resources for self-benefit rather than helping others. However, this self-serving bias may not extend to the point where it generates strong negative feelings in others which could result in significant feelings of guilt and risk of punishment for social norm violations.

## Acknowledgments

This work was supported by the National Natural Foundation of China (NSFC) grant numbers 31530032 and 91632117.

## Author contributions

X.X., B.B, and K.M.K designed the study and wrote the paper. X.X., C.L., Y.C. carried out the study. X.X., X.Z., Z.G., F.Z., and J.K. analyzed the data. All authors contributed to and have approved the final version of the manuscript.

## Supplementary information

In order to examine if there was any differential effect in the PLAY session of the Cyberball game between the first 4 blocks where participants did not receive a monetary reward and last 2 blocks where they did, we included the monetary condition as an independent variable in the analysis. Given the high correlation between ball tosses to excluder and victim players in the two stages (excluder player r = 0.477, *p* < 0.001; victim player r = 0.340, *p* = 0.002), we used the ball tosses to new player as a baseline and the dependent variable was used by subtracting ball tosses to new player from that of the excluder and victim players. A 2 (Treatment: PLC/OXT) × 2 (Player: Bad/Victim) × 2 (Monetary condition: First 4/Last 2) repeated ANOVA was performed and results showed there was no significant effect of monetary condition (main effect: *p* = 0.154; interaction: monetary condition × drug *p* = 0.420; player × monetary condition *p* = 0.409; player × monetary condition × drug *p* = 0.146) which indicated that subjects were consistent in their pattern of play during the entire PLAY session.

